# Liver-directed AAV gene therapy metabolically corrects AKU in *Hgd* deficient mice

**DOI:** 10.64898/2026.06.01.729174

**Authors:** Sien Lequeue, Brendan P. Norman, Gigly G. Del’haye, Jessie Neuckermans, Haaike Colemonts-Vroninks, Juliette H. Hughes, Matthias Rombaut, Paul Claes, Anja Heymans, Yves Heremans, Gunter Leuckx, Anneleen Mortier, Lakshminarayan Ranganath, James A. Gallagher, Tamara Vanhaecke, George Bou-Gharios, Joery De Kock

## Abstract

**Background:** Alkaptonuria (AKU) is a rare autosomal recessive metabolic disorder caused by deficiency of homogentisate 1,2-dioxygenase (HGD), resulting in systemic accumulation of homogentisic acid (HGA), ochronosis, and progressive multisystem disease. Although nitisinone (NTBC) lowers HGA levels, it does not correct the underlying genetic defect and induces hypertyrosinemia, highlighting the need for curative treatment approaches. We evaluated liver-directed adeno-associated virus (AAV)-mediated HGD gene therapy as a potential treatment for AKU.

**Methods:** Hgd-deficient (*Hgd*^*-/-*^) mice received liver-directed AAV2/8 vectors expressing codon-optimized human HGD under a liver-specific promoter. Reporter vectors were first used to assess hepatic biodistribution and transduction efficiency. Therapeutic efficacy was subsequently evaluated following AAV2/8-HGD administration (1 x 10^12^ vg/mouse). HGD expression was assessed by DNAscope, Western blotting, and RT-qPCR. Metabolic correction was determined using targeted LC-MS/MS and untargeted LC-HRMS metabolomics and compared with NTBC-treated *Hgd*^*-/-*^ mice.

**Results:** Reporter studies demonstrated liver-predominant transduction, with dose-dependent hepatocyte transduction reaching 89-93% at the highest dose. AAV2/8-HGD treatment produced robust hepatic HGD expression, with codon-optimized human HGD transcript levels approximately 33-fold higher than endogenous murine *Hgd* expression. Twelve weeks after treatment, plasma and urinary HGA levels were significantly reduced, with plasma HGA restored to near wild-type concentrations. Untargeted metabolomics further demonstrated marked reductions in HGA-derived phase I and II metabolites and revealed significant modulation of tricarboxylic acid cycle metabolism, consistent with partial restoration of metabolic homeostasis. Compared with NTBC-treated mice, AAV2/8-HGD achieved comparable plasma HGA reduction without elevation of upstream tyrosine pathway metabolites.

**Conclusions:** Liver-directed AAV2/8-HGD gene therapy achieved substantial biochemical correction in *Hgd*^*-/-*^ mice and restored metabolic flux without inducing hypertyrosinemia. These findings provide proof-of-concept supporting AAV-mediated HGD replacement as a promising long-term therapeutic strategy for AKU.

---

Alkaptonuria (AKU) is a rare autosomal recessive disorder caused by biallelic mutations in homogentisate 1,2-dioxygenase (HGD), an enzyme primarily expressed in liver and kidney, leading to systemic accumulation of homogentisic acid (HGA) (Fig. S1). Excess HGA undergoes oxidation and polymerization into a melanin-like pigment that deposits in connective tissues (ochronosis), causing progressive early-onset arthropathy and multiple systemic manifestations. These include pigmentation of the skin, eyes, and ears, as well as cardiovascular and renal complications such as aortic stenosis, vascular calcification, and kidney stone formation. While nitisinone (NTBC), the current standard of care, reduces HGA by inhibiting 4-hydroxyphenylpyruvate dioxygenase (HPD), it causes hypertyrosinemia and does not correct the underlying genetic defect, and therefore must be combined with a lifelong protein-restricted diet, underscoring the need for a curative therapy (Fig. S1).^1^ Liver-directed AAV gene therapy is a promising approach for monogenic liver diseases like AKU, but has not yet been tested in this context. Evidence from a liver-specific AKU mouse model and a clinical liver transplant case indicates that the liver is the primary site of HGA clearance, and partial restoration (26-43% of normal) of hepatic HGD is sufficient to normalize plasma HGA levels, whose elevation drives ochronosis in AKU.^2^ These findings support the evaluation of liver-directed AAV-HGD therapy as a potential treatment for AKU. We therefore evaluated liver-directed AAV2/8 gene therapy delivering codon-optimized human HGD under a liver-specific promoter in the *Hgd tm1a*^*-/-*^ (*Hgd*^*-/-*^) mouse model, which carries a gene-trap disrupting *Hgd* expression, closely recapitulating elevated HGA and joint pigmentation.^2^ Therefore, our study aimed to determine whether liver-directed AAV2/8-mediated HGD expression can restore biochemical correction in AKU, providing proof-of-concept for a gene therapy approach.

To determine liver-specific biodistribution and transduction, eight-week-old *Hgd*^*-/-*^ males were injected with the AAV2/8-fLUC-P2A-GFP reporter vector (Fig. S2A) at doses of 1 x 10^12^, 0.5 x 10^12^, or 1 x 10^11^ vg/mouse, with the experimental timeline detailed in Fig. 1A. Bioluminescence imaging (BLI) at 1, 2, and 4 weeks post-injection showed liver-predominant expression, with photon emission visibly decreasing at lower vector doses (Fig. 1B and Fig. S3). DNAscope *in situ* hybridization at four weeks showed dose-dependent liver transduction efficiencies of 89.3% (±7.5%), 40.85% (±1.6%), and 24.13% for 1 x 10^12^, 0.5 x 10^12^, or 1 x 10^11^ vg/mouse, respectively (Fig. 1B and Fig. 1C).

**Figure 1.**
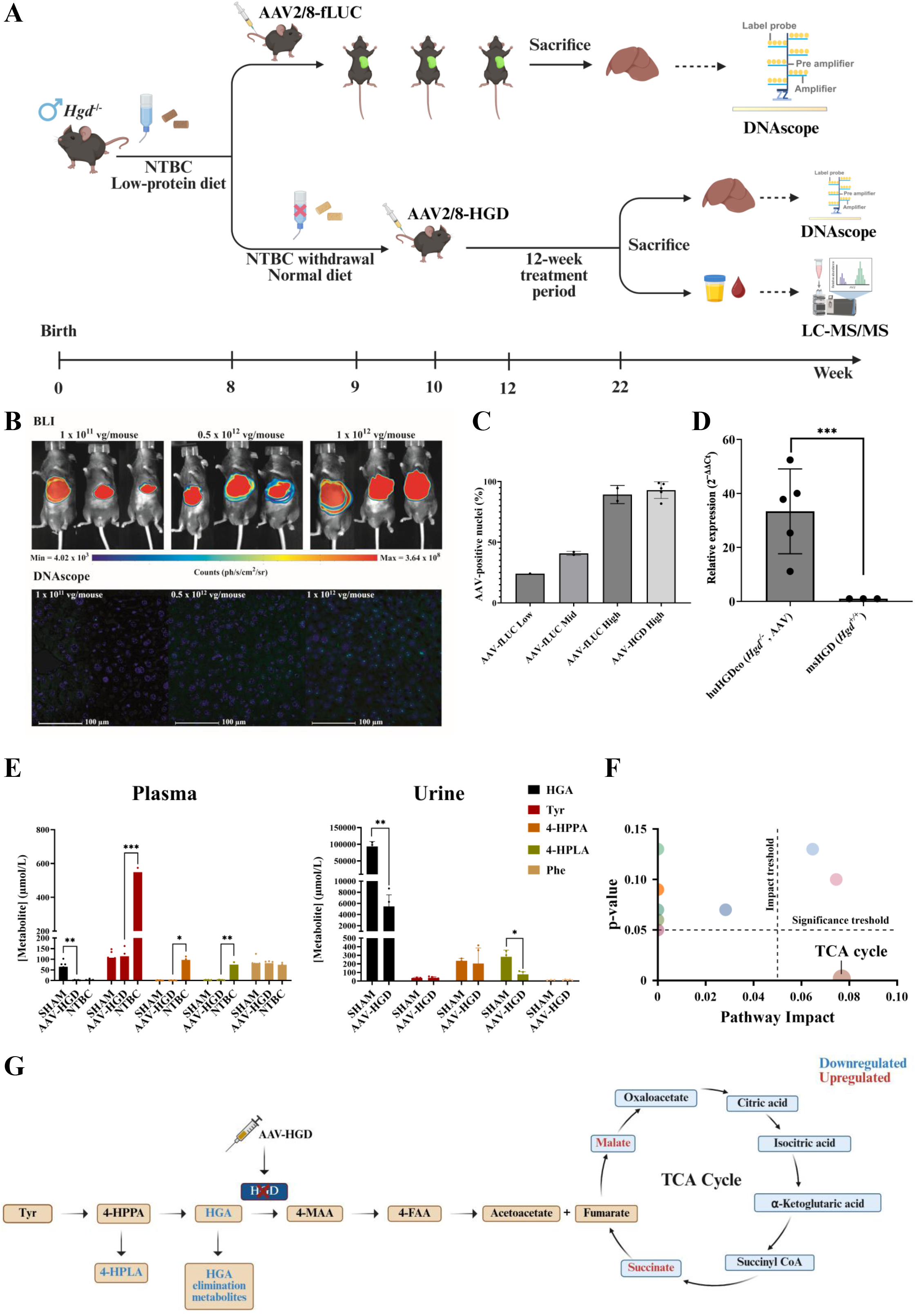
Experimental design and *in vivo* outcomes of liver-directed AAV2/8-HGD gene therapy in *Hgd*□*/*□ mice. (A) Overview of *in vivo* experimental designs. All mice were eight weeks old at study initiation and received continuous NTBC treatment (4 mg/L) in drinking water and a low Tyr/Phe diet *ad libitum* unless otherwise specified, to reduce circulating HGA levels and mitigate NTBC-induced hypertyrosinemia. Upper panel: Experiment 1 – Biodistribution study. Male *Hgd*^-/-^ mice received AAV2/8-fLUC-P2A-GFP at 1 x 10^11^ (N = 1), 0.5 x 10^12^ (N = 2), or 1 x 10^12^ (N = 2) vg/mouse. Bioluminescence imaging (BLI) was performed at 1, 2, and 4 weeks post-injection to assess vector distribution and expression. At week 4, mice were sacrificed and livers were collected for DNAscope *in situ* hybridization to quantify successfully transduced hepatocytes. Lower panel: Experiment 2 – Therapeutic study. After a 2-week NTBC washout on a standard fat and protein diet, *Hgd*^□^*/*^□^ mice received either AAV2/8-HGD (N = 5) or sterile saline (N = 6). Mice remained off NTBC throughout the experiment. Urine was collected at week 12 post-injection, followed by sacrifice, plasma collection by cardiac puncture, and liver isolation. Plasma and urine metabolites were analysed using LC-QQQ-MS and LC-QTOF-MS, and hepatocyte transduction efficiency was assessed in liver sections by DNAscope *in situ* hybridization. (B) BLI and liver transduction efficiencies following AAV2/8-fLUC-P2A-GFP administration. Upper panel: Representative BLI at 4 weeks post-injection for all three AAV2/8-fLUC-P2A-GFP doses, illustrating liver-predominant expression with visibly lower photon emission at reduced vector doses, while higher doses produced saturated signals that limited qualitative assessment of liver specificity. The photon emission range was 4.02 x 10^3^ (blue) to 3.64 x 10^8^ photons per second per square centimeter per steradian (ph/s/cm^2^/sr) (red). Additional images from the same doses, captured at one- and two-weeks post-injection, are provided in Figure S3 for comparison. Lower panel: DNAscope *in situ* hybridization images of liver sections at 4 weeks post-injection showing dose-dependent AAV probe signal (green) colocalized with DAPI-stained nuclei (blue) in hepatocytes. (C) Quantification of hepatocyte transduction efficiencies by DNAscope. Liver transduction was quantified by identifying membrane-segmented hepatocytes positive for both AAV probe signal and a DAPI-stained nucleus. For AAV2/8-fLUC-P2A-GFP, transduction efficiencies at four weeks were 24.13% at the low dose (1 x 10^11^ vg/mouse, N = 1), 40.85% ± 1.6% at the mid dose (5 x 10^11^ vg/mouse, N = 2), and 89.3% ± 7.5% at the high dose (1 x 10^12^ vg/mouse, N = 2). For AAV2/8-HGD, the transduction efficiency at week 12 post-injection was 92.9% ± 6.9% (N = 5). (D) Relative HGD expression in AAV-treated mice. Bar plots show relative expression (2^-ΔΔCt^) of codon-optimized human HGD (huHGDco) in AAV-treated *Hgd*^□^*/*^□^ livers compared to endogenous mouse HGD (msHGD) in WT *Hgd*^*+/+*^ livers (WT = 1). Normality of ΔCt values was assessed using the Shapiro-Wilk test, and an unpaired t-test with Welch’s correction was performed (AAV huHGDco, N = 5; WT MsHGD, N = 3; p < 0.001). Data are presented as mean ± SD. huHGDco expression in AAV-treated livers was strongly elevated compared to endogenous msHGD, with a mean fold increase of 33.3 ± 15.7. (E) Targeted metabolic profiling of Tyr pathway metabolites. Plasma and urine metabolites were measured in AAV2/8-HGD-treated *Hgd*^□^*/*^□^ mice and compared to SHAM controls; plasma metabolites were also compared to a separate NTBC-treated cohort (N = 3) maintained on a high-fat, low-protein diet as a standard-of-care reference (Table S1 and Table S4). Data are presented as mean ± SD for normally distributed metabolites and median (25^th^-75^th^ percentile) for non-normally distributed metabolites. Normality was assessed by Shapiro-Wilk test; statistical significance was determined using Welch’s t-test (normally distributed) or Mann-Whitney test (non-normal plasma HPPA), with p ≤ 0.05 considered significant (* p ≤ 0.05, ** p ≤ 0.01, *** p ≤ 0.001). In AAV2/8-HGD-treated mice, plasma and urine HGA were significantly reduced 35.4- and 17.1-fold, respectively (p = 0.002 and p = 0.008), and urine 4-hydroxyphenyllactate (4-HPLA) decreased 3.6-fold (p = 0.04), while Phe, Tyr, and 4-hydroxyphenylpyruvate (4-HPPA) remained unchanged. Comparison to NTBC-treated mice showed significant reductions in plasma 4-HPLA (p = 0.006), 4-HPPA (p = 0.04), and Tyr (p < 0.001), whereas HGA and Phe were similar between groups. Note: Urine data for the 12-week SHAM group and plasma data for the 12-week NTBC group are based on three biological replicates due to limited sample availability. (F) Pathway analysis of altered metabolites following AAV2/8-HGD gene therapy in *Hgd*^*-*/-^ mice. Untargeted metabolomics was performed on plasma and urine samples to identify metabolic differences between the 12-week SHAM and 12-week AAV2/8-HGD-treated groups (Table S2). These metabolites were then analysed using MetaboAnalyst 6.0 to determine modulated pathways (Table S3). The bubble chart provides a visual summary of the top 10 pathways identified by MetaboAnalyst analysis, showing their relative importance following treatment. The x-axis represents pathway impact, reflecting the relative importance of each pathway, while the y-axis displays p-value, indicating statistical significance. Notably, the TCA cycle was the only pathway that exceeded the predefined pathway impact threshold (≥ 0.05) and met the significance criterion (p ≤ 0.05), indicating it was significantly modulated following gene therapy. (G) Modulated metabolic pathways in AAV2/8-HGD treated *Hgd*^-/-^ mice. The figure illustrates the modulation of key metabolic pathways, including the Tyr degradation pathway and the TCA cycle following treatment with AAV2/8-HGD. Metabolites within each pathway are shown with their respective upregulation (red) or downregulation (blue) after treatment. These pathways were selected based on previous studies linking them to the AKU phenotype^4^. Data are presented to highlight the metabolic alterations observed in response to the gene therapy. Abbreviations: Tyr: Tyrosine, 4-HPPA: 4-hydroxyphenylpyruvate, 4-HPLA: 4-hydroxyphenyllactate, 4-MAA: 4-maleylacetoacetate, 4-FAA: 4-fumarylacetoacetate.

We next evaluated the therapeutic AAV2/8-HGD vector (Fig. S2B) at the highest dose (1 x 10^12^ vg/mouse), confirming comparable liver transduction (92.9% ± 6.9%; Fig 1C and Fig. S4), with the experimental timeline shown in Fig. 1A. Western blot (Fig. S5) showed a 100 kDa HGD band, possibly a multimeric form, in AAV2/8-HGD-treated *Hgd*^*-/-*^ and wildtype (WT, *Hgd*^*+/+*^) mice, indicating successful expression, and absent in PBS-treated *Hgd*^*-/-*^ mice as expected. RT-qPCR analysis demonstrated that, in AAV2/8-HGD-treated *Hgd*^□^*/*^□^ mice, the relative expression of codon-optimized human HGD (huHGDco) was markedly increased compared with endogenous mouse *Hgd* expression (msHGD) in WT (*Hgd*^*+/+*^) mice, with a mean (SD) fold increase of 33.3x (±15.7) (p□< □0.001), indicating strong transgene overexpression (Fig. 1D). Subsequently, metabolic correction was assessed by targeted LC-MS/MS and untargeted LC-HRMS. Twelve weeks post-injection, targeted LC-MS/MS analysis of protein-depleted EDTA plasma and urine showed that SHAM-treated mice maintained high HGA levels (65.2□±□26.0□μmol/L and 93□641□±□14□575□μmol/L, respectively), consistent with disease baseline.^2^ In AAV2/8-HGD-treated mice, plasma and urine HGA were significantly reduced 35.4- and 17.1-fold, respectively (p = 0.002 and p = 0.008). Urine 4-hydroxyphenyllactate (4-HPLA) also decreased significantly (3.6-fold, p = 0.04), while phenylalanine (Phe), tyrosine (Tyr), and 4-hydroxyphenylpyruvate (4-HPPA) levels remained unchanged in both plasma and urine (Fig. 1E, Table S1). Untargeted metabolomics analysis was performed on plasma and urine from 12-week AAV2/8-HGD-treated and SHAM control mice. Altered metabolites were identified according to predefined thresholds (see Methods and Table S2). Significant reductions were observed in HGA phase I and II biotransformation products, including plasma HGA-glucuronide (FC = 11.2, q = 0.008) and urine HGA-hydroxysulfate (FC = 6.6, q = 0.007), HGA-demethylation/hydroxylation products (FC = 5.7, q = 0.04), HGA-glucuronide (FC = 26.4, q = 0.08), and acetyl-HGA (FC = 1.6, q = 0.06) (Table S2). In addition, metabolomics data were analysed using MetaboAnalyst, evaluating 10 pathways with significance thresholds of p ≤ 0.05 and impact ≥ 0.05. Only the citrate (TCA) cycle was significantly altered (p = 0.002, impact = 0.08) (Table S3, Fig. 1F & 1G).

To benchmark AAV2/8-HGD efficacy against the current standard of care, targeted LC-MS/MS was performed on plasma from a separate cohort of *Hgd*^*-/-*^ mice treated for 12 weeks with NTBC combined with a high-fat, low-protein diet and compared to 12-week AAV2/8-HGD-treated mice. Analytes included HGA, 4-HPPA, 4-HPLA, Tyr, and Phe (Fig. 1E and Table S4). Plasma analysis showed significant reductions in 4-HPLA (p = 0.006), 4-HPPA (p = 0.04), and Tyr (p < 0.001) in AAV2/8-HGD-treated mice compared to NTBC-treated mice, while plasma HGA and Phe were similar between groups.

Taken together, high-dose AAV2/8-HGD gene therapy restored plasma HGA in *Hgd*^-/-^ mice to near-WT levels (1.7□± □0.6□μmol/L), whose elevation drives ochronosis, the key pathophysiological event in AKU.^1–3^ Biodistribution and DNAscope *in situ* hybridization analyses confirmed efficient hepatic transduction (92.9%), and RT-qPCR showed huHGDco expression ∼33-fold higher than endogenous msHGD. Although complete liver correction is not required, as reported in literature (26-43% of normal hepatic HGD mRNA), the observed hepatocyte transduction and the high huHGDco expression contributed to full metabolic rescue, consistent with normalization of circulating HGA. However, our data indicate that urine HGA levels were not fully normalized. Although high liver transduction strongly reduced urinary HGA, complete correction was not achieved, likely reflecting the lack of targeted kidney HGD expression. This is consistent with literature showing that kidney HGD primarily influences urinary, but not plasma, HGA levels.^2^ Further metabolic profiling revealed significant reductions in HGA phase II biotransformation products, including sulfated and glucuronidated conjugates, in treated mice compared to controls (Table S2 and Fig. 1G). This reflects a shift from compensatory phase II metabolism, normally upregulated in AKU to detoxify excess HGA, toward enhanced degradation through the canonical Tyr pathway, restoring metabolic flux toward a healthy phenotype.^4^ Moreover, in AAV2/8-HGD-treated mice, 4-HPLA levels were significantly reduced, indicating partial correction of upstream Tyr metabolism, which aligns with previous findings in *Hgd*^*-/-*^ mice showing altered Tyr, Phe, and pheripheral neurotransmitter metabolism, likely driven by elevated HGA, which appears partially reversed by gene therapy. Pathway analysis further revealed partial restoration of the TCA cycle, including malic and succinic acid, consistent with literature suggesting that HGD deficiency disrupts fumarate availability and energy metabolism in AKU.^4^ Correcting hepatic HGD with AAV2/8-HGD thus not only normalizes HGA but also helps restore energy metabolism. Importantly, comparison with NTBC-treated *Hgd*^*-/-*^ mice showed similar reductions in plasma HGA, but unlike NTBC, AAV2/8-HGD did not elevate upstream Tyr metabolites such as Tyr, 4-HPLA, or 4-HPPA. NTBC blocks HPD upstream, causing hypertyrosinemia^1^, whereas AAV2/8-HGD corrects the underlying hepatic HGD deficiency without introducing a metabolic bottleneck, providing a more physiological restoration of Tyr metabolism.

In conclusion, these results provide proof-of-concept that AAV2/8-HGD gene therapy can achieve biochemical correction in AKU. Correction was achieved with ∼93% hepatocyte transduction, but efficiency is expected to be lower in larger models.^5^ Future studies could explore whether lower, human-relevant doses can achieve similar metabolic correction. Unlike NTBC, gene therapy restores hepatic HGD without upstream metabolite accumulation, supporting liver-directed AAV-HGD as a potential long-term treatment for AKU.

## Data availability statement

The authors confirm that the data supporting the findings and conclusion of this study are presented within the article and its supplemental information.

## Acknowledgements

This study is financially supported by the Research Foundation Flanders (FWO) under grant number G023320N (EvolvAKUre). Figures created with BioRender.com with their individual figure URLs are as follows: Figure 1A: https://BioRender.com/oe0hudb, Figure 1G: https://BioRender.com/or8enx0, Figure S2A: Created in https://BioRender.com/r48g777, Figure S2B: https://BioRender.com/r33p050. We would like to thank Prof. Dr. George Bou-Gharios and Dr. Brendan Norman from the University of Liverpool (UoL) from the Department of Musculoskeletal and Ageing Science for providing their mouse model and their expertise in metabolomics. We also extend our gratitude to Dr. Yves Heremans from the Visual and Spatial Tissue Analysis (VSTA) core facility (https://vsta.research.vub.be) and Gunter Leuckx from the VUB from the Laboratory of Beta Cell Neogenesis, for their expertise in DNAscope *in situ* hybridization and quantification, respectively.

## CRediT authorship contribution statement

Conceptualization, S.L. and J.D.K.; Investigation, S.L., B.N., G.D.H., J.N., H.C.-V., P.C., A.H., Y.H., G.L.; Methodology, S.L., B.N., G.D.H., J.N., H.C.-V., Y.H., A.M., J.H.H., J.D.K. Resources, J.D.K., T.V., J.A.G. and G.B.-G. Data Curation, S.L. and J.D.K.; Writing – Original Draft Preparation, S.L. and J.D.K.; Writing – Review and Editing, S.L., B.N., G.D.H., J.N., H.C.-V., J.H.H., M.R., Y.H., A.M., L.R., J.A.G., T.V., G.B.-G. and J.D.K.; Visualization, S.L. and J.D.K.; Supervision, T.V., L.R., J.A.G., G.B.-G. and J.D.K. Funding Acquisition, T.V., J.A.G., G.B.-G. and J.D.K. All authors have read and agreed to the published version of the manuscript.

## Ethics declaration

The animal study was approved by the Institutional Animal Ethics Committees of Vrije Universiteit Brussel (VUB), Belgium (approval numbers 20-210-3, 20-210-9, and 21-210-2).

## Declaration of Generative AI and AI-assisted technologies in the writing process

During the preparation of this work, the authors used ChatGPT to improve the clarity and readability of certain sentences by refining grammar and restructuring complex phrases. After using this tool, the authors reviewed and edited the content as needed and take full responsibility for the content of the publication.

