## Supplementary Data for "Liver-directed AAV gene therapy metabolically corrects AKU in *Hgd* deficient mice"

**Materials and Methods**

***Hgd*^-/-^ mouse model and housing**

Experiments were performed using the well-characterized AKU mouse model *Hgd* *tm1a^-/-^* (*Hgd*^-/-^) (C57BL/6 background), in which a gene trap cassette containing an IRES:LacZ and promoter-driven neo cassette is inserted between exons five and six, thereby disrupting endogenous *Hgd* expression and generating a constitutive global knockout. In homozygous *Hgd^-/-^* mice, ochronosis resembles the early stages of human joint disease, with pigment primarily confined to individual chondrocytes and their surrounding matrix in the calcified cartilage.^1^ While the model can be used to assess joint pathology, in this study we focused exclusively on evaluating systemic interventions for restoring HGD activity and correcting the associated metabolic abnormalities. This constitutive knockout model differs from the conditional models *Hgd* *tm1c* (wildtype, WT) and *Hgd* *tm1d* (liver-specific *Hgd* deletion), which are generated by Flp- or Cre-mediated recombination to remove the gene trap cassette and allow tissue-specific or inducible control of *Hgd* expression.^1^ Mouse care, breeding, and experiments were approved by the Institutional Animal Ethics Committees (under grant numbers 20-210-3, 20-210-9, 21-210-2) and carried out at Vrije Universiteit Brussel (VUB), Belgium. Mice were group-housed pathogen-free in individually ventilated cages with a 14/10 light/dark cycle, 19-23°C, and 30-70% relative humidity. All mice received continuous NTBC treatment (4 mg/L) *via* their drinking water and were fed a low Tyr and Phe diet (LabDiet® 5LJ5 chow, LabDiet, St. Louis, MO, USA) *ad libitum* unless specified otherwise, to reduce circulating HGA levels and mitigate NTBC-induced hypertyrosinemia, consistent with clinical management of AKU. We have made every effort to report the results in accordance with the ARRIVE guidelines.^2^

**Genotyping**

Genotyping was performed by extracting genomic DNA tissue from an ear punch using the REDExtract-N-Amp Tissue PCR kit (Merck KGaA, Darmstadt, Germany) following the manufacturer’s instructions. Polymerase chain reaction (PCR) amplification was carried out according to the protocol by Hughes *et al*.^1^, followed by electrophoresis (RT, 100 V, 300 mA, 1 h) on a 2% *(w/v)* agarose gel (SERVA Electrophoresis Gmbh) stained with Gelred^®^ (Biotum, Brussels, Belgium). Visualization and fragment size determination were conducted using ChemiDoc MP (Bio-rad, Temse, Belgium) and a 100 base pair (bp) Generuler (Fisher Scientific, Waltham, MA, USA). Mutants exhibited two modified alleles located at approximately 257 bp, whereas WT (two unmodified alleles) showed the 561 bp band and heterozygotes (one modified allele and one unmodified allele) showed the 561 bp and 257 bp bands.

**AAV vector engineering and production**

A liver-specific thyroxine-binding globulin (TBG) promoter was used to ensure strong, liver-restricted transgene expression. To maximize hepatic transduction, we selected the AAV8 capsid combined with an AAV2 backbone, providing high liver tropism and efficient vector production.^3,4^ AAV vectors encoding fLUC-P2A-GFP (Genbank: PQ897897) and codon-optimized WT human HGD for *Mus musculus* (Genbank: PQ897898) (NM_000187.3) were generated with a liver-specific TBG promoter. The transgene cassette was inserted into an AAV2 transfer plasmid and confirmed by Sanger sequencing (Eurofins Genomics, Ebersberg, Germany). High-titer AAV2/8 vectors were produced and purified by SignaGen Laboratories (Frederick, MD, USA).

**Biodistribution and liver transduction efficiency of** **AAV2/8-fLUC-P2A-GFP in *Hgd*^-/-^ mice using bioluminescence imaging and DNAscope**

Eight-week-old *Hgd*^-/-^ male mice (28.8 ± 2.0 g) were intravenously injected with three doses of AAV2/8-fLUC-P2A-GFP to determine liver transduction efficiencies: 1 x 10^12^ vg/mouse (N = 2), 0.5 x 10^12^ vg/mouse (N = 2), 1 x 10^11^ vg/mouse (N = 1). Although a bicistronic construct encoding fLUC and GFP was generated, only fLUC expression was used for downstream biodistribution and *in vivo* imaging experiments. BLI was performed on all injected mice (three per group), irrespective of injection success, to visualize liver-specific transgene expression, whereas DNAscope quantification of liver transduction efficiency was limited to mice that received the full intended dose. BLI was performed at one, two- and four-weeks post-injection using a PhotonIMAGER^TM^ Optima System (Biospace, Nesles La Vallée, France). Mice were anesthetized with 2% isoflurane and oxygen and were intraperitoneally injected with D-luciferin (150 µg/g body weight) (Promega, Madison, WI, USA) for imaging. At four weeks post-injection, mice were sacrificed, and liver samples were collected and washed with ice-cold PBS to remove any residual debris. Tissue pieces measuring approximately 1-2 cm^3^ were fixed in 4% *(w/v)* paraformaldehyde (Sigma-Aldrich) at 4°C for 24 hours. Following fixation, liver tissue samples were dehydrated through a graded ethanol series (VWR, Leuven, Belgium), embedded in paraffin using a Microm GmbH STP 120-1 tissue processor (Prosan, Arnhem, The Netherlands), and sectioned for histological analysis. DNAscope *in situ* hybridization was then performed on these liver sections using the RNAscope Multiplex Fluorescent v2 Assay kit (Cat. No. 323100, Advanced Cell Diagnostics, Newark, CA, USA), according to the manufacturer’s instructions with adapted pre-treatment conditions (15 min. Target Retrieval, 15 min. Protease Plus diluted 1/5 in PBS). Probes used were pAAV-TBG-promoter-sense (Cat No. 114958-C1) and *Mus musculus* Ppib (Cat No. 313911-C2), detected with Vivid 520 and Vivid 570 at 1/1000 (Bio-Techne, Minneapolis, MN, USA), respectively. Imaging was performed using an Axio Scan.Z1 (Zeiss, Jena, Germany), and quantitative spatial analysis was conducted in HALO 3.2 (Indica Labs, Albuquerque, NM, USA). For each image, a single region of interest (ROI) was selected. Hepatocyte identification and counting were performed using the HALO AI membrane segmentation algorithm, which segments cells based on membrane boundaries. As a result, multinucleated hepatocytes were counted as individual cells. Nuclear detection algorithms assessed DAPI-stained nuclei, which were used to identify nuclei within each membrane-segmented cell. A cell was considered DAPI-positive if at least one DAPI-stained nucleus was detected inside the segmented membrane boundary. AAV-probe positivity was determined by detecting AAV signal colocalized with DAPI-stained nuclei within the same segmented cell. The percentage of positive cells was calculated relative to the total number of membrane-segmented cells in each ROI. DNAscope *in situ* hybridization was subsequently performed on liver tissues from AAV2/8-HGD injected *Hgd*^-/-^ mice (N = 5) to verify liver transduction efficiencies of the AAV2/8-HGD construct.

**Evaluation of therapeutic efficacy of hepatotropic AAV2/8-HGD**

Eight-week-old *Hgd*^-/-^ male mice (29.0 ± 1.5 g) were withdrawn from NTBC two weeks prior to AAV administration and switched from a high-fat and low-protein diet (LabDiet^®^ 5LJ5 chow) to a standard fat and protein diet (Safe 105^®^, Safe Diets, Augy, France). Male mice were used because AKU, though affecting both sexes equally, manifests earlier and with greater severity in this sex, and limiting the study to males reduces hormonal variability while modeling a more pronounced biochemical phenotype.^5^ After this two-week washout period, mice were injected intravenously (100 µL, tail vein) using a 30Gx^1/2^” needle (0.3 x 12 mm; BBraun, Machelen, Belgium) with either sterile PBS (SHAM control, N = 6) or AAV2/8-HGD (treatment, N = 5). Mice remained on the standard diet without NTBC for the remainder of the study. Urine samples were collected 12 weeks post-injection for metabolic profiling. At the end of the study (also 12 weeks post-injection), mice were anesthetized by intraperitoneal injection of a ketamine (87.5 mg/kg Ketamidor^®^)/xylazine (12.5 mg/kg Rompun^®^) mixture. Blood was collected by ventral heart puncture using a 26Gx^1/2”^ needle (0.45 x 13 mm) and a 1 mL syringe (Terumo^®^, VWR, Leuven, Belgium). Approximately 0.5-1 mL of blood was drawn into EDTA-coated microtubes (Sarstedt K3E tubes, Nümbrecht, Germany) and centrifuged at 1500 x *g* for 15 minutes at 4°C. Plasma fractions were then isolated and stored at -80°C for metabolic profiling. In addition, plasma from a separate cohort of 12-week NTBC-treated *Hgd^⁻/⁻^* mice (N=3), maintained on a high-fat and low-protein diet, was included for benchmarking against the standard of care. For protein and gene expression studies, liver tissue sections covering the entire liver were collected (total tissue volume up to 1 cm^3^), preserved in RNAprotect Tissue Reagent (Qiagen, Hilden, Germany), and stored at -80°C until further analysis. Acidified plasma^6,7^ and urine^7,8^ samples were measured for concentrations of Tyr pathway metabolites using published LC-QQQ-MS assays. The metabolites measured were Phe, Tyr, 4-HPPA, 4-HPLA, and HGA. Untargeted metabolic profiling of plasma and urine samples was conducted following a previously established LC-QTOF-MS protocol.^9^

**Quantitative real-time PCR**

Total RNA was extracted using the RNeasy Mini Kit (Qiagen) according to the manufacturer’s instructions. Reverse transcription was performed using the iScript™ cDNA Synthesis Kit (Bio-Rad). Quantitative PCR (qPCR) was conducted using the PowerUp™ SYBR™ Green Master Mix (Thermo Scientific) following the manufacturer’s instructions, on a QuantStudio^TM^ 3 Real-Time PCR system (Thermo Scientific). Gene-specific primers were designed to assess expression of the codon-optimized human HGD (HuHGDco) delivered *via* AAV and the endogenous mouse *Hgd* (msHGD) to determine the relative proportion of vector-derived HGD mRNA. Several candidate mouse housekeeping genes were evaluated for stability using the BestKeeper algorithm, and mouse actin beta (msACTB) and mouse TATA-box binding protein (msTBP) were selected for normalization. The sequences of the primers used were as follows: HuHGDco forward 5’ - GCTTCGGCAACGAGTGCAGCA - 3’, reverse 5’ - TACAGCCAG CTTCTCTTGTTG - 3’; msHGD forward 5’ - GATTTGGGAATGAGTGTGCTT - 3’, reverse 5’ - TACAGC CAGCTTCTCTTATTG - 3’. Housekeeping genes msACTB forward 5’ - GGCTGTATTCCCCTCCATCG - 3’, reverse 5’ - CCAGTTGGTAACAATGCCATGT - 3’; msTBP forward 5’ CCTTGTACCCTTCACCAATGAC 3’, reverse 5’ - ACAGCCAAGATTCACGGTAGA - 3’. qPCR was performed on RNA extracted from multiple liver sections spanning the entire liver, and cycle threshold (Ct) values were averaged to obtain a single value per liver. Two housekeeping genes, msACTB and msTBP, were used for normalization, and the geometric mean of their Ct values was calculated for each mouse. Target gene expression of both msHGD and vector-derived huHGDco was normalized to this geometripc mean by calculating ΔCt values, defined as the difference between the target gene Ct and the average housekeeping gene Ct within the same mouse. To compare expression across groups, the geometric mean of ΔCt values for msHGD in three WT livers was calculated. ΔΔCt values were then determined by subtracting this mean ΔCt of WT msHGD from the ΔCt of huHGDco in AAV-treated livers. Relative expression levels were calculated using the 2^−ΔΔCt^ method, which assumes comparable and near-optimal amplification efficiencies for all primer pairs. All primer pairs generated single specific products as confirmed by melt-curve analysis. Bar plots show relative expression (2^−ΔΔCt^) of HuHGDco in AAV-treated livers compared to msHGD in WT livers (WT = 1). Normality of ΔCt values for both groups was assessed using the Shapiro-Wilk test. Subsequently, an unpaired t-test with Welch’s correction was performed on the ΔCt values to compare AAV HuHGDco and WT msHGD.

**Western blot**

Frozen liver tissues from *Hgd*^-/-^ mice injected with AAV2/8-HGD, sterile saline (negative control), and C57BL/6 WT mice (positive control) were homogenized and protein concentrations were determined using a bicinchoninic acid BCA assay.^10^ For sodium dodecyl sulfate-polyacrylamide gel electrophoresis and Western blot analysis, 50 µg of protein was separated on a 12% *(w/v)* Mini-Protean^®^ TGX Stain-Free™ gel (Bio-Rad) and transferred onto a nitrocellulose membrane using Trans-Blot^®^ Turbo™ transfer packs (Bio-Rad). Membranes were blocked for one hour with 5% *(m/v)* milk powder in Tris-buffered saline (20 mM Tris, 135 mM NaCl) supplemented with 0.1% *(v/v)* Tween-20, followed by overnight incubation at 4°C with a polyclonal rabbit anti-HGD antibody (ProteinTech, Rosemont, IL, USA; 1:10 000 in blocking buffer). After washing, membranes were incubated for one hour at room temperature with a polyclonal goat anti-rabbit secondary antibody (Dako, Agilent Technologies, Heverlee, Belgium; 1:1 000 in blocking buffer). Protein bands were visualized using the Pierce™ ECL Western Blotting Substrate Kit (Thermo Scientific) and imaged with the ChemiDoc MP Imaging System (Bio-Rad).

**Data processing and statistical analysis**

Data obtained from targeted and untargeted metabolomic assays were further subjected to statistical evaluation and pathway analysis using the following tools and criteria. For targeted metabolomics, we used Graphpad Prism 9.5.1 software (Graphpad Software Inc, San Diego, CA, USA). The data were expressed as means with SD. Normal distribution was confirmed using the Shapiro-Wilk test. We performed a two-sample, two-tailed Welch’s t-test for 12-week AAV2/8-HGD vs. 12-week SHAM samples. For data that were not normally distributed (plasma 4-HPLA levels), the Mann-Whitney test was applied, and these data are summarized as median with interquartile ranges (25^th^ - 75th percentile). Significance was determined based on probability (p) values, with values ≤ 0.05 considered statistically significant. A post-hoc power analysis was performed using G*Power v3.1 (Heinrich Heine University, Düsseldorf, Germany) to determine whether the study had sufficient power to detect the observed difference in plasma HGA, the main metabolite responsible for AKU pathology. The observed means, SDs, and sample sizes were used to calculate Cohen’s d. Power was then calculated using a two-sample, two-tailed t-test with an alpha of 0.05. For plasma HGA, levels in AAV2/8-HGD-treated mice (1.84 ± 1.13 µmol/L, N = 5) compared to SHAM controls (65.16 ± 25.98 µmol/L, N = 6) gave a Cohen’s d of ~3.26, corresponding to a post-hoc power of 0.99. This confirms that the sample size was sufficient to detect a significant difference. For untargeted metabolomics, we utilized Progenesis QI v2.3 software (Nonlinear Dynamics, Newcastle, UK). This software aligned retention time (RT), performed peak picking (also filtering out noise and background signals) and deconvolution, normalized the data and can finally identify metabolites. Metabolite characteristics were determined using a specialized database for accurate mass retention time (AMRT) constructed under identical analytical conditions. The database included AMRT targets from IROA Technology MS metabolite standards^11^ and phase I and II biotransformation products relevant to the Phe-Tyr pathway metabolites^12^ associated with AKU. Semi-targeted metabolite identification was conducted using the Human Metabolome Database (HMDB 2022, v5)^13^ and the Biotransformer database (BioTransformer Rails application 2022, version 3.1.0)^14,15^, allowing ion species (M-H^-^) and (M+H^+^) in negative and positive polarity, respectively, with an accuracy range of ± 5 ppm for accurate mass (AM) only. Metabolites meeting the criteria of q-value ≤ 0.1, FC ≥ ±1.2, coefficient of variation ≤ 30%, and abundance greater than 1000 were selected for further analysis using MetaboAnalyst 6.0^16^ for pathway analysis. A q-value threshold of 0.1 was chosen to increase sensitivity, allowing detection of biologically relevant Tyr-Phe pathway metabolites that would have been missed with a more stringent cutoff, while still controlling the false discovery rate. The mouse-specific KEGG pathway library (*Mus musculus*) was used as the reference metabolome. Pathway analysis was performed using the hypergeometric test, and pathway topology was analysed by relative-betweenness centrality. Visualization of results was done through scatter plots showing all pathways, with reference lines indicating the significance threshold (p ≤ 0.05) and pathway impact threshold (≥ 0.05). Metabolites lacking KEGG or HMDB identifiers, including 4-hydroxyphenyllactate (4-HPLA), glutamyl-Tyr, N-acetyl-L-serine, and several HGA phase II biotransformation products (acetyl-HGA, HGA-glucuronide, HGA-hydroxysulfate, HGA-demethylation/hydroxylation), were excluded from the pathway analysis. Due to the limited number of metabolites included, pathway coverage and robustness may be affected, and findings should be interpreted with caution.

**Supplementary Figures**

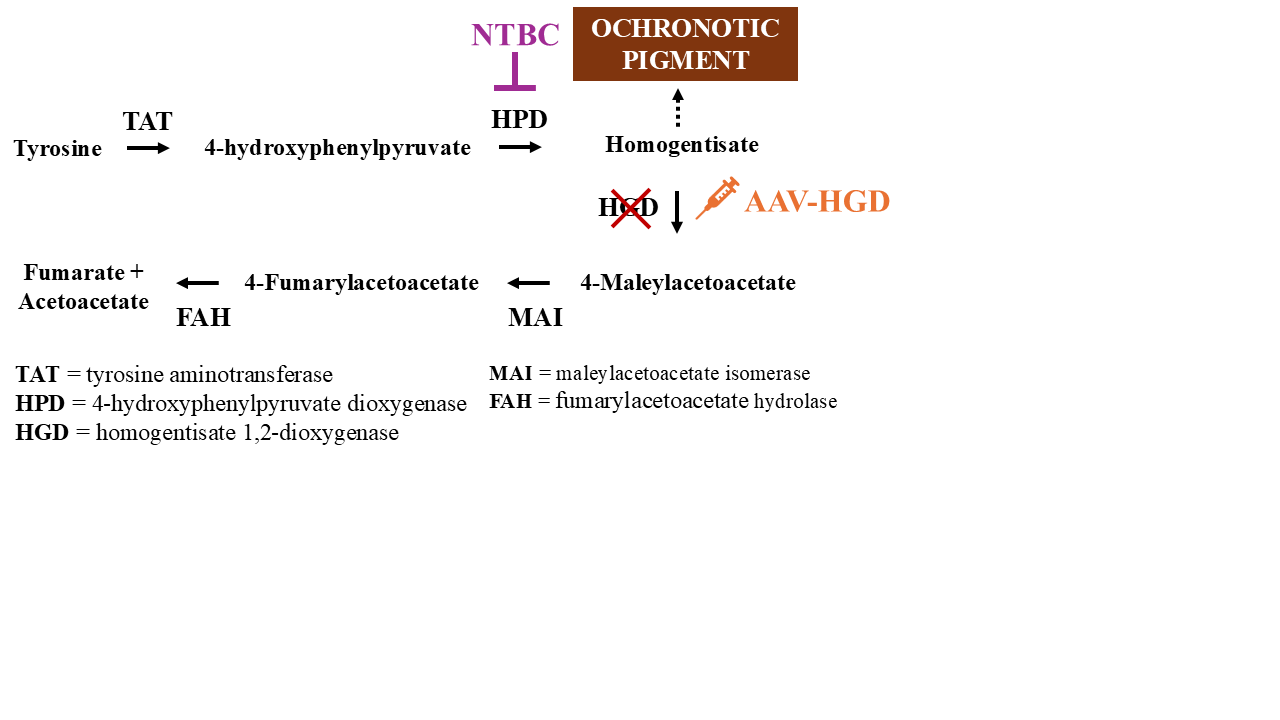

**Supplementary Figure 1.** Schematic representation of the Tyr degradation pathway highlighting the role of HGD in converting HGA into MAA. HGD deficiency leads to the accumulation of HGA, resulting in AKU, with NTBC being the only current treatment option. The figure additionally highlights the site of action of the AAV-HGD construct, intended to restore functional HGD activity.

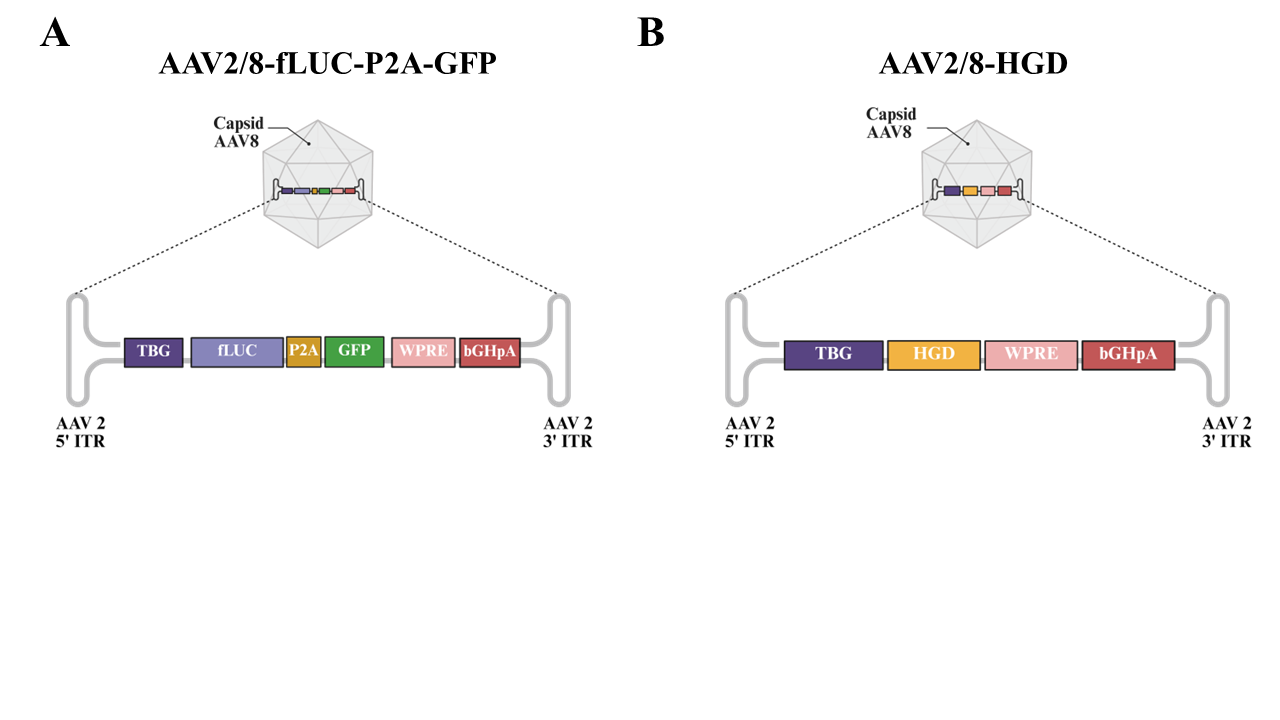

**Supplementary Figure 2. AAV2/8 vector constructs used in this study**. (A) AAV2/8-fLUC-P2A-GFP and (B) AAV2/8-HGD both contain the liver-specific thyroxine-binding globulin (TBG) promoter and the woodchuck hepatitis virus post-transcriptional regulatory element (WPRE) to enhance transgene expression.

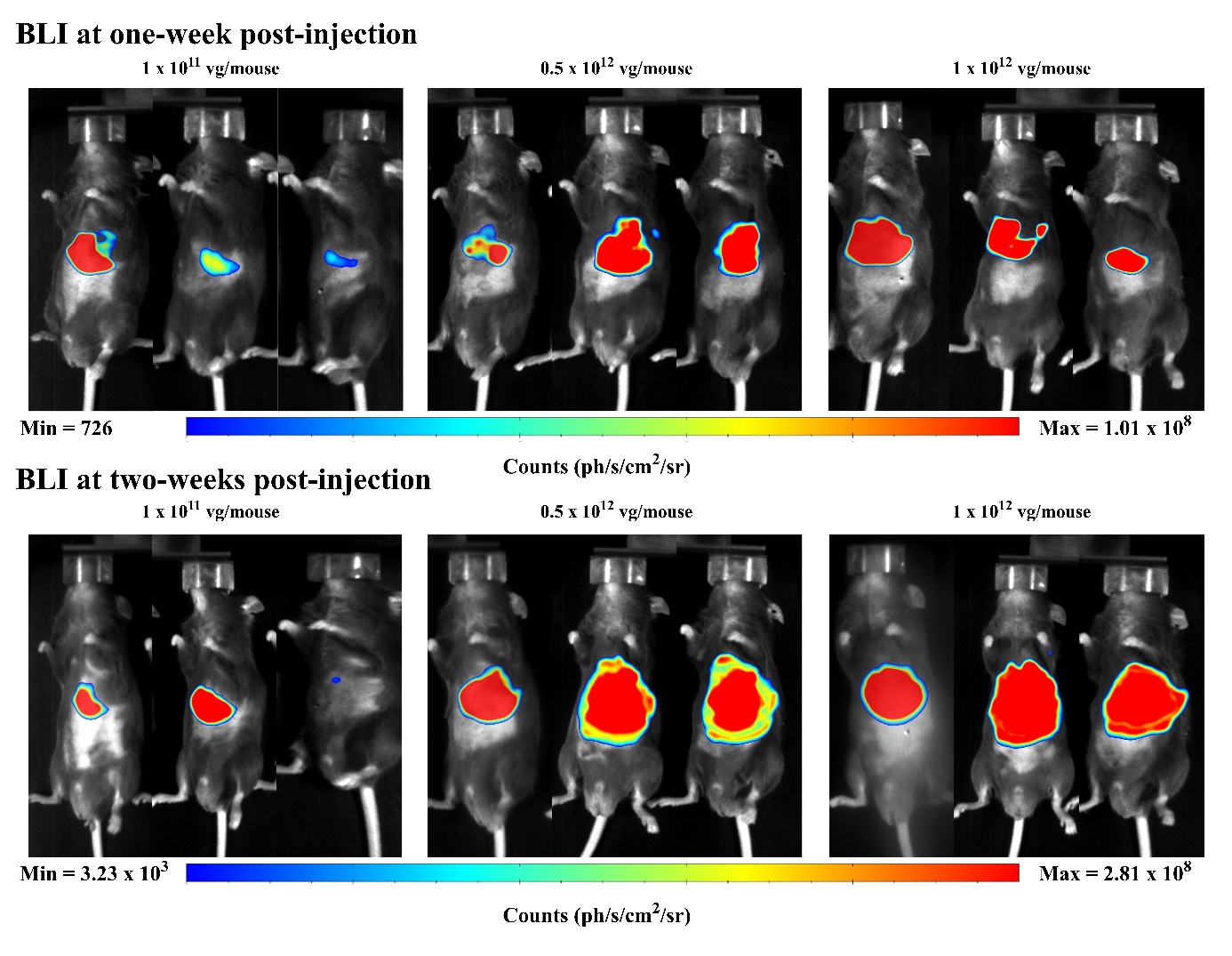
**Supplementary Figure 3. Evaluation and confirmation of AAV2/8-fLUC-P2A-GFP vector construct.** *Hgd^-/-^* mice were intravenously injected with three different doses of AAV2/8-fLUC-P2A-GFP (1 x 10^11^ vg/mouse, 0.5 x 10^12^ vg/mouse, and 1 x 10^12^ vg/mouse). BLI was performed one- and two-weeks post-injection. **The** photon emission range for one-week post-injection was **726 (blue) to 1.01 x 10^8^** photons per second per square centimeter per steradian **(ph/s/cm²/sr) (red)**, and for **two-weeks post-injection**, it ranged from **3.23 x 10^3^ (blue) to 2.81 x 10^8^ ph/s/cm²/sr (red)**. Signal was predominantly localized to the liver, visibly decreasing at lower vector doses. At higher doses, signal intensity was elevated, limiting qualitative assessment of liver specificity. Additional images from the same doses, captured at four weeks post-injection, are provided in **Figure 1B**.

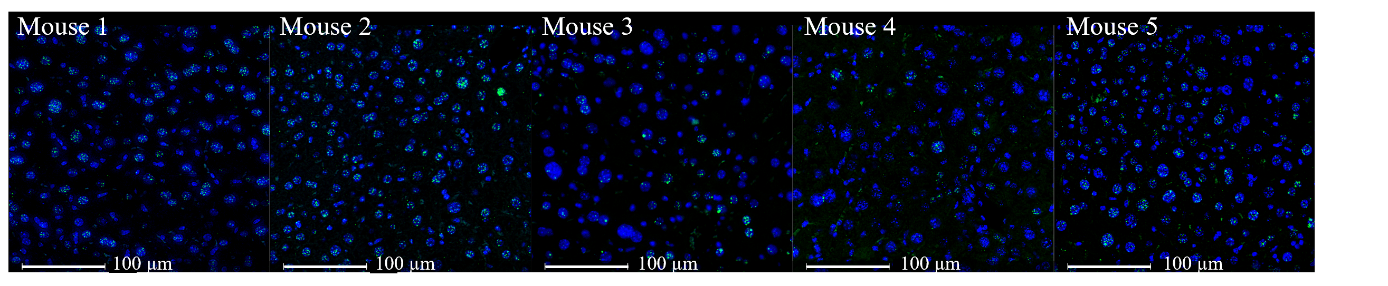

**Supplementary Figure 4. Liver transduction of AAV2/8-HGD in *Hgd^-/-^* mice.** DNAscope *in situ* hybridization was performed on liver tissues from *Hgd^⁻/⁻^* mice injected with AAV2/8-HGD (1 x 10^12^ vg/mouse, N = 5). Representative images show AAV probe signal (green) colocalized with DAPI-stained nuclei (blue) in hepatocytes from each injected mouse. Scale bar = 100 µm. Quantification of transduction efficiencies is shown in Figure 1C.

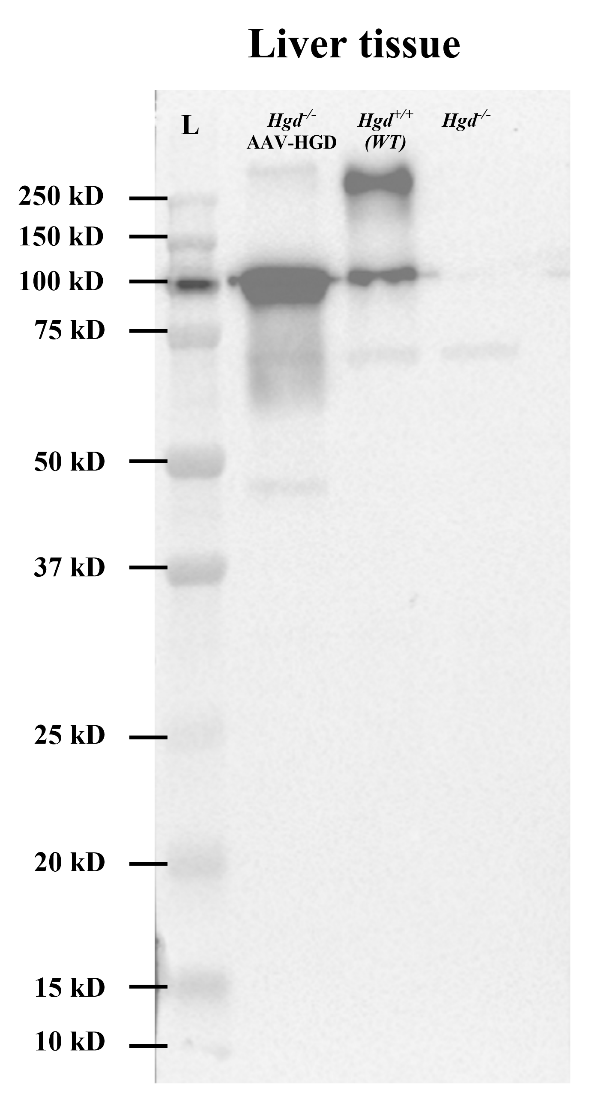

**Supplementary Figure 5.** Western blot of liver homogenates shows a distinct band at approximately 100 kDa in *Hgd^-/-^* mice injected with AAV-HGD and WT (*Hgd^+/+^*) mice, but not in saline-injected controls. This indicates the successful expression of HGD in the liver following AAV-HGD injection. The band at 100 kDa likely represents a multimeric form of the HGD protein. Additionally, a protein band is observed at a higher molecular weight (>250 kDa) in both AAV-HGD injected *Hgd^-/-^* mice and *Hgd^+/+^*mice, which possibly corresponds to the hexameric form of the HGD protein. As these bands are also present in WT mice and absent in negative controls, this suggests they represent forms of endogenous HGD.

**Supplementary Tables**

**Supplementary Table 1. Summary of urinary and plasma Tyr pathway metabolite levels (µmol/L) in treated AAV2/8-HGD and untreated (SHAM) *Hgd^-/-^* mice, analysed using a targeted metabolomics approach.** Data are presented as mean ± SD for normally distributed metabolites and as median with interquartile range (25^th^-75^th^ percentile) for non-normally distributed metabolites. Normality was assessed using the Shapiro-Wilk test. A two-sample, two-tailed Welch’s t-test was used for normally distributed data, while the Mann-Whitney test was applied to non-normally distributed plasma HPPA levels. P-values ≤ 0.05 were considered statistically significant.

| **Metabolite** | **Biofluid** | **AAV2/8-HGD (µmol/L)** | **SHAM (µmol/L)** | **p-value** |
| --- | --- | --- | --- | --- |
| Phe | Plasma | 81.82 (± 10.99) | 80.62 (± 28.02) | 0.93 |
|  | Urine | 13.34 (± 4.46) | 11.57 (± 2.57) | 0.50 |
| Tyr | Plasma | 114.10 (± 31.82) | 107.70 (± 33.37) | 0.75 |
|  | Urine | 37.93 (± 12.16) | 35.69 (± 6.88) | 0.75 |
| HGA | Plasma | 1.84 (± 1.13) | 65.16 (± 25.98) | 0.002 |
|  | Urine | 5 473.51 (± 2 068.77) | 93 641.22 (± 14 575.89) | 0.008 |
| 4-HPPA | Plasma | 0.19 (0.00 - 1.38) | 0.00 (0.00 - 1.33) | 0.59 |
|  | Urine | 203.90 (± 181.50) | 235.20 (± 27.99) | 0.72 |
| 4-HPLA | Plasma | 2.56 (± 0.83) | 2.08 (± 1.04) | 0.41 |
|  | Urine | 78.54 (± 29.99) | 282.41 (± 77.82) | 0.04 |

**Supplementary Table 2. Summary of significantly altered metabolites detected in plasma and urine of AAV2/8-HGD treated mice when compared to untreated control mice.** Identified metabolites were listed with their respective biofluid in which they were detected, Human Metabolome Database (HMDB) code, mass-to-charge ratio (m/z), retention time (RT), fold change (FC), false discovery rate-adjusted p-value (q-value), the basis for their compound identification and polarity (electrospray ionization, ESI). Metabolite identification was performed based on accurate mass and retention time (AMRT), the matching criteria were ± 10 ppm and ± 0.3 min, respectively, and matched against a database created internally using metabolite standards. Metabolite identification performed on accurate mass (AM) alone was conducted with a mass tolerance of ± 5 ppm and matched against both the HMDB database and the BioTransformer database. Metabolites are ordered by their FC with those highlighted in red indicating a positive FC and those in blue indicating a negative FC.

| **Metabolite** | **Biofluid** | **HMDB** | **m/z** | **RT (min.)** | **FC** | **q-value** | **compound ID** | **ESI** |
| --- | --- | --- | --- | --- | --- | --- | --- | --- |
| HGA-glucuronide | Urine | NA | 343,068 | 2,63 | -26,41 | 0,08 | AMRT | NEG |
| HGA-glucuronide | Plasma | NA | 343,067 | 2,60 | -11,16 | 0,008 | AMRT | NEG |
| HGA-hydroxysulfate | Urine | NA | 262,989 | 2,97 | -6,63 | 0,007 | AMRT | NEG |
| HGA-demethylation  and hydroxylation | Urine | NA | 169,015 | 3,61 | -5,70 | 0,04 | AMRT | NEG |
| Glutamyl-Tyr | Plasma | HMDB0028831 | 311,125 | 3,71 | -4,13 | 0,098 | AMRT | POS |
| Biliverdin IX | Plasma | HMDB0001008 | 581,241 | 11,51 | -2,52 | 0,05 | AM | NEG |
| N-Acetyl-L-serine | Urine | HMDB0002931 | 146,046 | 1,64 | -2,19 | 0,04 | AMRT | NEG |
| Acetyl-HGA | Urine | NA | 209,046 | 6,47 | -1,61 | 0,06 | AMRT | NEG |
| Indolepyruvate | Plasma | HMDB0060484 | 202,051 | 5,97 | 2,04 | 0,008 | AM | NEG |
| Succinic acid | Urine | HMDB0000254 | 117,020 | 2,33 | 4,32 | 0,06 | AMRT | NEG |
| Malic acid | Urine | HMDB0000156 | 133,015 | 1,55 | 8,53 | 0,03 | AMRT | NEG |

**Supplementary Table 3.** Pathway Analysis performed by MetaboAnalyst. The table provides an overview of each analysed metabolic pathway, including the total number of metabolites, the number of significantly altered metabolites, the p-value, the pathway impact, and the altered metabolites. Pathways were considered significant at p ≤ 0.05 and a pathway impact ≥ 0.05. For a visual summary of the top 10 pathways identified by MetaboAnalyst, showing their relative importance following treatment, see Figure 1F in the main article.

| **Pathway Name** | **Total metabolites** | **Hits** | **p-value** | **Pathway impact** | **Altered metabolites** |
| --- | --- | --- | --- | --- | --- |
| Citrate cycle (TCA cycle) | 20 | 2 | 0.002 | 0.08 | Malic acid, succinic acid |
| Butanoate metabolism | 15 | 1 | 0.05 | 0.0 | Succinic acid |
| Ubiquinone and other terpenoid-quinone biosynthesis | 19 | 1 | 0.06 | 0.0 | Homogentisic acid |
| Propanoate metabolism | 22 | 1 | 0.07 | 0.0 | Succinic acid |
| Pyruvate metabolism | 23 | 1 | 0.07 | 0.03 | Malic acid |
| Alanine, aspartate and glutamate metabolism | 28 | 1 | 0.09 | 0.0 | Succinic acid |
| Porphyrin metabolism | 31 | 1 | 0.10 | 0.08 | Biliverdin |
| Glyoxylate and dicarboxylate metabolism | 32 | 1 | 0.13 | 0.0 | Malic acid |
| Tryptophan metabolism | 41 | 1 | 0.13 | 0.0 | Indolepyruvate |
| Tyrosine metabolism | 42 | 1 | 0.13 | 0.07 | Homogentisic acid |

**Supplementary Table 4. Summary of plasma Tyr pathway metabolite levels (µmol/L) in treated AAV2/8-HGD and NTBC *Hgd^-/-^* mice, analysed using a targeted metabolomics approach.** Data are presented as mean ± SD for normally distributed metabolites and as median with interquartile range (25^th^-75^th^ percentile) for non-normally distributed metabolites. Normality was assessed using the Shapiro-Wilk test. A two-sample, two-tailed Welch’s t-test was used for normally distributed data, while the Mann-Whitney test was applied to non-normally distributed HPPA levels. P-values ≤ 0.05 were considered statistically significant.

| **Metabolite** | **AAV2/8-HGD (µmol/L)** | **NTBC (µmol/L)** | **p-value** |
| --- | --- | --- | --- |
| Phe | 81.82 (± 10.99) | 74.46 (± 9.41) | 0.36 |
| Tyr | 114.10 (± 31.82) | 548.5 (± 23.01) | < 0.001 |
| HGA | 1.84 (± 1.13) | 4.88 (± 3.51) | 0.27 |
| 4-HPPA | 0.19 (0.00 - 1.38) | 96.82 (± 18.72) | 0.04 |
| 4-HPLA | 2.56 (± 0.83) | 74.99 (± 9.87) | 0.006 |
